# Effects of chemical modulators on enzyme specificity

**DOI:** 10.1101/2025.09.30.679615

**Authors:** Andrew D. Hecht, Oleg A. Igoshin

## Abstract

Chemical inhibitors bind to enzymes, thereby inhibiting their catalytic activity. While many enzymes catalyze reactions with a single substrate, others, like DNA polymerase, can act on multiple related substrates. Substrate-selective inhibitors (SSIs) target these multi-substrate enzymes to modulate their specificity. Although SSIs hold promise as therapeutics, our theoretical understanding of how different inhibitors influence enzyme specificity remains limited. In this study, we examine enzyme selectivity within kinetic networks corresponding to known inhibition mechanisms. We demonstrate that competitive and uncompetitive inhibitors do not affect substrate specificity, regardless of rate constants. In contrast, noncompetitive and mixed inhibition can alter specificity and can lead to non-monotonic responses to the inhibitor. We show that mixed and non-competitive inhibitors achieve substrate-selective inhibition by altering the effective free-energy barriers of product formation pathways that are enabled by the inhibitor’s presence. We then apply this framework to the Sirtuin-family deacylase SIRT2, showing that the suicide inhibitor thiomyristoyl lysine (TM) cannot influence substrate specificity unless there is a direct substrate exchange reaction or biochemical constraints are relaxed. These findings provide insights into engineering systems where cofactor binding modulates metabolic flux ratios.

## Introduction

Enzymes are proteins that catalyze chemical reactions involving one or more substrates, thereby playing key roles in cellular function. Given the importance of enzymes in biology, many pharmaceutical drugs target enzymatic processes to produce medically beneficial effects; these drugs are often chemical inhibitors that can slow or halt the activity of a target enzyme. For example, nonsteroidal anti-inflammatory drugs (NSAIDs) act as inhibitors of the cyclooxygenases COX-1 and COX-2 and reduce the production of pro-inflammatory prostaglandins.^1^ Tyrosine kinase inhibitors^2,3^ and histone deacetylase inhibitors^4^ have been used as cancer therapies to promote apoptosis in cancer cells.

Many enzymes can be post-translationally regulated by small-molecule inhibitors; these inhibitors can be divided into three major classes based on their interaction with the free enzyme and the enzyme-substrate complex. In the case of competitive inhibitors, the inhibitor molecule directly competes with substrates for access to the enzyme active site. ^5^ In contrast, uncompetitive inhibitors do not interfere with substrate binding and instead bind to the enzyme-substrate complex. ^5^ Finally, non-competitive inhibitors (and the related mixed inhibitors) are capable of binding to both the free enzyme and enzyme-substrate complex. ^5^ The mechanism of a particular inhibitor is biochemically relevant as it governs the precise manner in which the kinetics of the inhibited enzyme are affected.

While many enzymes act on only a single substrate, in some cases, enzymes such as DNA polymerase can catalyze reactions involving various chemically similar substrates. For these multi-substrate enzymes, each unique substrate can lead to a corresponding product. Importantly, an enzyme is not necessarily equally efficient with all substrates; depending on the enzyme’s ability to discriminate between substrates, some products will be more prevalent than others even with equal substrate concentrations.

Inhibitors that can affect the substrate selectivity of the targeted enzyme, known as substrate-selective enzyme inhibitors (SSIs), are becoming increasingly important and represent a novel class of drugs. In one example, Maianti et al. carried out a high-throughput screen to identify SSIs capable of preferentially inhibiting insulin degradation by insulin-degrading enzyme (IDE) without affecting glucagon degradation.^6^ As insulin and glucagon have opposite effects on blood glucose levels, SSIs targeting IDE could serve as a new class of drug for treating type II diabetes. SSIs are also relevant to basic research, as potential confounding factors for reverse chemical genetics screens. ^7^

Kinetic modeling can provide insights into how enzyme inhibition affects substrate specificity. Previously, Hendricks et al. developed a kinetic model of the kinase p38 to explain substrate-selective inhibition of substrate phosphorylation. ^8^ However, to our knowledge, there has been no comprehensive theoretical study on the effects of different enzyme inhibition modalities on substrate specificity.

Prior work by Mallory et al. ^9^ demonstrated that kinetic barrier perturbations are required to modulate the steady-state ratio of reaction fluxes for alternative biochemical pathway branches. On the other hand, merely changing the stability of states is insufficient to affect the flux ratio.^9^ These findings imply that inhibitors that operate by effectively stabilizing certain states will not be substrate-selective.

Here, we employ kinetic models of common inhibition modalities for multi-substrate enzymes to examine the possible effects of inhibition on substrate selectivity. We demonstrate that competitive and uncompetitive inhibitors can never affect substrate selectivity, regardless of the underlying kinetic parameters governing the system. In contrast, mixed and partial inhibition modes can not only affect substrate selectivity, but can also yield non-monotonic effects where the substrate selection error can have a local maximum/minimum for intermediate inhibitor concentrations. We examine the thermodynamic basis for substrate-selective inhibition by mixed and partial inhibitors, and show that substrate selectivity can only change when the presence of the inhibitor leads to an alternative product formation pathway with potentially different kinetic barriers. Finally, we construct a kinetic model of substrate selection by the Sirtuin-family de-acylase SIRT2 and show that either direct substrate exchange reactions or a relaxed, unordered binding mechanism are required to observe substrate-selective inhibition.

## Theoretical Methods

### Kinetic Models of Enzyme Inhibition Modes

Reversible enzyme inhibitors are primarily classified by the inhibition mechanism, which is determined by the structure of both the enzyme and the inhibitor molecule. The most basic form of inhibition is competitive inhibition (Figure 1A), in which the inhibitor directly competes with substrate for access to the enzyme active site. ^5^ This competition effectively raises the enzyme Michaelis-Menten constant, *K*_M_, i.e., higher substrate concentrations are required to achieve the same reaction velocity. In contrast to competitive inhibitors that bind to the free enzyme, uncompetitive inhibitors bind directly to the enzyme-substrate complex (Fig. 1B) and block the catalytic transformation. Uncompetitive inhibitors generally reduce the maximum enzyme velocity, *V*_max_, but can also affect the observed *K*_M_,.^5^

**Figure 1.**
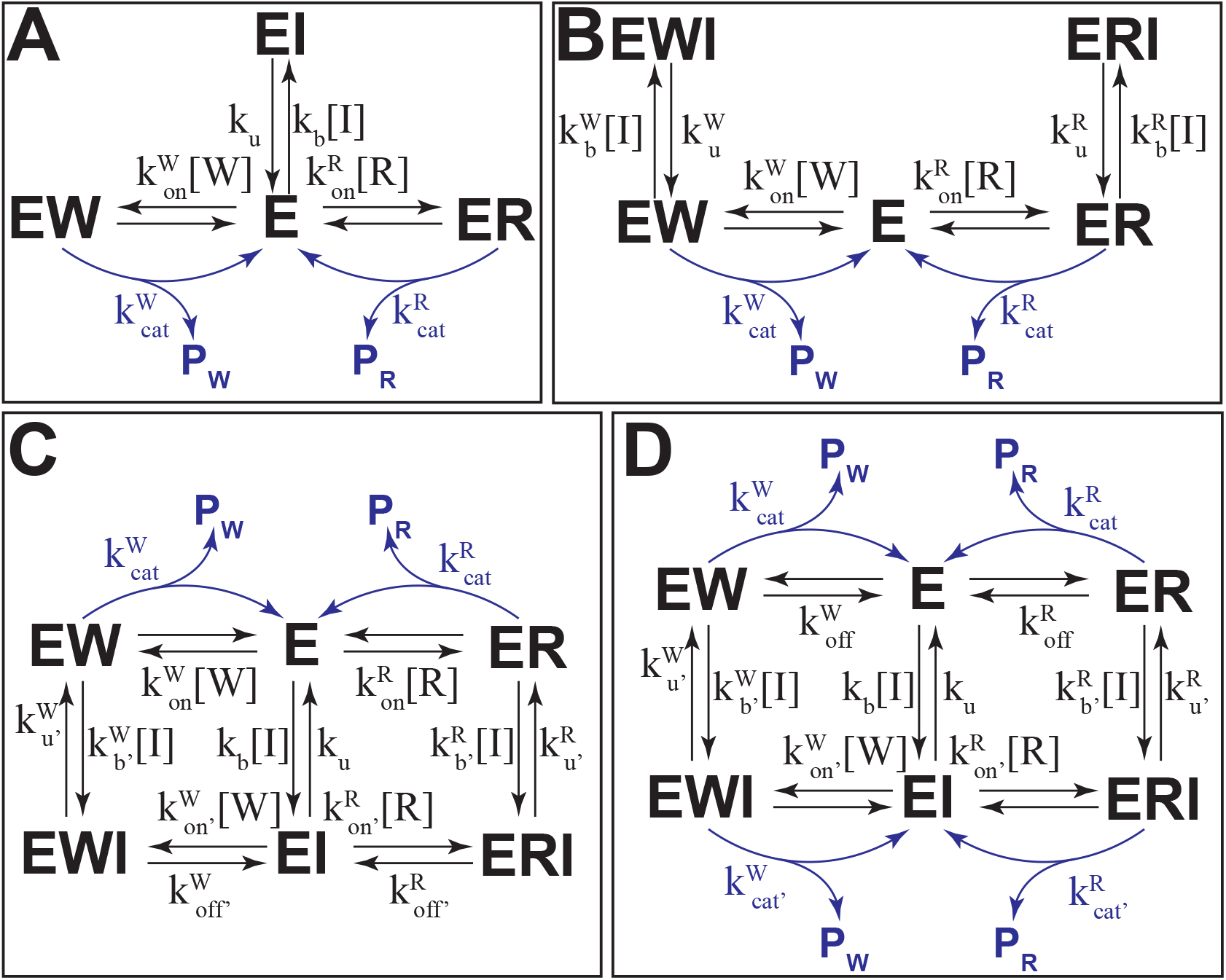
Kinetic schemes for competitive (A), uncompetitive (B), mixed (C), and partial inhibition (D). Product formation reactions are shown in blue.

Mixed inhibitors combine features from competitive and uncompetitive inhibition, and can bind to both the free enzyme and the enzyme-substrate complex (Figure 1). If the inhibitor binds equally well to both enzyme states, the inhibitor is said to be a non-competitive inhibitor, which is a special case of mixed inhibition. Both mixed and non-competitive inhibitors can affect the enzyme *V*_max_, however, only mixed inhibition can affect the apparent *K*_M_.^5^ Finally, inhibition is occasionally incomplete, leading to partial inhibition (Fig. 1D). In this case, residual catalytic activity can be observed even when fully bound to inhibitor. ^10^

To examine the effects of each inhibition mode on substrate selectivity, we construct kinetic models for each of the schemes in Figure 1; the models are built using techniques previously used to study substrate selection errors in various biological systems, ^11–13^ and allow us to examine the behavior of each inhibition mode analytically. While in principle, enzymes can have a large number of possible substrates, we limit ourselves to two substrates, R and W, for simplicity.

The forward chemical master equation allows us to compute the likelihood of observing each state in the model, and is defined by a system of ordinary differential equations for each fixed value of inhibitor concentration, *x*:

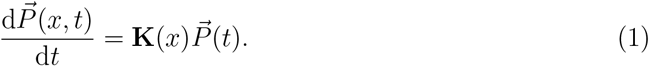

with the normalization constraint

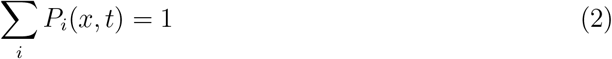

The matrix **K** encodes the state transitions for each system, with **K**_*ji*_ = *k*_*ij*_ for each transition *i* → *j*. The rate constants for the R and W substrates are related through discrimination factors, which are defined as

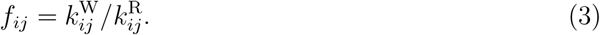

Additionally, we assumed substrate, cofactor, and inhibitor concentrations are kept constant and absorbed these into the rate constants. We only keep the dependence on the inhibitor concentration (*x*) explicit. A detailed description of each of the inhibition models is provided in the Supplemental Information.

The steady-state probabilities 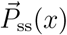 can be computed by enforcing the additional constraint

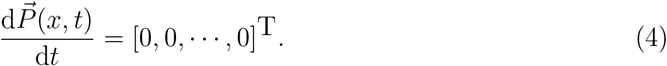

The fluxes for each reaction in the system can be readily computed from the system probabilities; for a given reaction *i* → *p*, the flux is simply *J*_*i*_ = *k*_*ip*_*P*_*i*_. The total product formation flux for each substrate can be defined as a sum over all possible product formation reactions; for a substrate with *N* such reactions, the total flux is simply

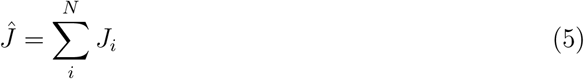

with the ratio of fluxes for two substrates, R and W, being given as

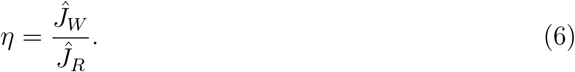

The flux ratio indicates the relative catalytic efficiency for the two substrates; as the flux ratio decreases, the enzyme increasingly favors substrate R over substrate W.

Kinetic schemes that contain thermodynamic (non-dissipative) reaction cycles, such as the mixed and partial inhibition models (Figure 1C,D), are additionally constrained by detailed balance. Detailed balance constrains the rate constants for reactions along the cycle such that the forward and reverse rates for each reaction *i* are related by

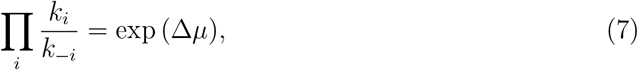

where *µ* is the chemical potential difference for the cycle; at equilibrium, Δ*µ* = 0 k_B_T.

### Parameter Sampling

To examine the prevalence of non-monotonic dose-response curves, model parameters were sampled in log-space using the MATLAB function `rand`. Given upper and lower bounds of *a* and *b*, for each *N* parameter model we computed a parameter set 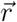 as

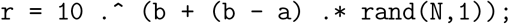

Generally, the rate constants are sampled from the range [10^−4^, 10^4^] for all models. The model discrimination factors are sampled separately, with the full bounds given in Supplemental Table 1.

For simplicity, we require that the saturating bound (*η*_∞_) be lower than the uninhibited bound (*η*_0_). In the case of mixed inhibition, this is satisfied for all parameter sets with *f*_p_ < *f*_off_; however, we were unable to derive a corresponding constraint for partial inhibition. Therefore, we enforced the constraint by rejecting parameter sets that fail to satisfy *η*_∞_ < *η*_0_. However, we still set the discrimination in catalysis, *f*_cat_ based on the sampled 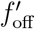 by scaling 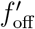 by a number sampled from the uniform distribution, i.e.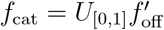. We also assume 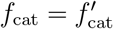 for partial inhibition.

### Detecting Undershoot and Overshoot

For each random parameter set, we first compute the uninhibited and saturation bounds *η*_0_ and *η*_∞_. Next, starting with an initial concentration range *x* ∈ [*x*_*i*_, *x*_*j*_], we compute the flux ratio *η*(*x*). In order to ensure that we capture the full range of behavior for each parameter set, we progressively expand the concentration range until the computed flux ratios are sufficiently close to the limits *η*_0_ and *η*_∞_.

Undershoot events occur when the flux ratio *η*(*x*) becomes smaller than the saturating limit *η*_∞_ for intermediate inhibitor concentrations. For a given parameter set, the total undershoot *q*_under_ is computed as the difference

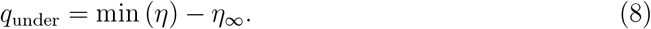

Similarly, we can compute the total overshoot *q*_over_ for a given parameter set as

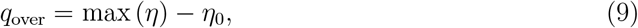

where we now compare the maximum flux ratio with the uninhibited limit.

In all cases, we enforce a threshold *ϵ* = 0.01 to avoid parameter sets that only display a slight undershoot or overshoot; when plotting, all parameter sets with undershoot or overshoot under the threshold *ϵ* are discarded.

## Results

### Competitive and Uncompetitive Inhibitors Cannot Affect Specificity

First, we determined how the flux ratio *η* changes as a function of inhibitor concentration *x* for each inhibition scheme (Figure 1, see theoretical methods for details). When the inhibitor is substrate-selective, we expect to see the flux ratio change with increasing inhibitor concentration, with the derivative 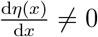 for at least some range of concentrations.

For competitive and uncompetitive inhibition (Figure 1A,B) we find that the concentration of inhibitor does not affect the product flux ratio (Figure 2A,B). Specifically, the flux ratio *η* is independent of the inhibitor concentration *x* for these inhibition modes and has the same form in both cases:

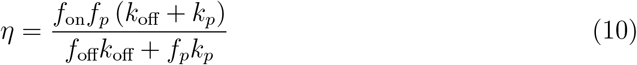

**Figure 2.**
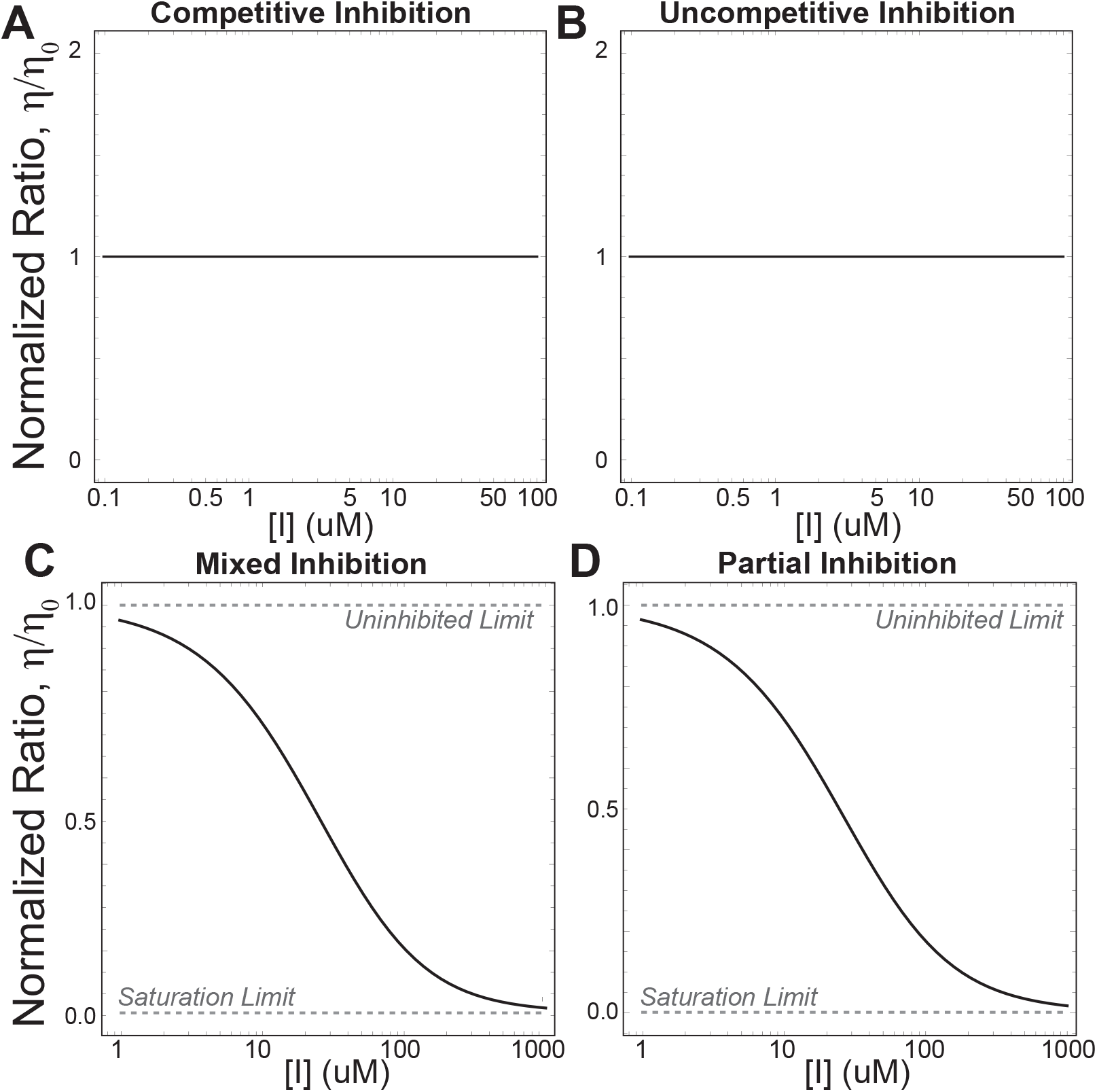
Competitive (A) and uncompetitive (B) inhibitors cannot affect substrate selectivity, no matter the underlying kinetic parameters. In contrast, both mixed (C) and partial inhibition (D) can yield substrate-selective inhibition.

Thus, regardless of kinetic parameters, the inhibitor cannot affect substrate selectivity for competitive or uncompetitive inhibition. Notably, an uncompetitive inhibitor that only inhibits one of the substrates (i.e., only binds to EW to form EWI but not to ER) is still incapable of affecting selectivity.

In contrast, both mixed inhibition (Figure 1C) and partial inhibition (Figure 1D) can affect substrate selectivity, and are therefore capable of being substrate-selective inhibitors (Figure 2C,D). In both cases, the structure of the reaction network involves reaction cycles (loops), which we assume are non-dissipative, i.e., their rate constants obey the detailed-balance constraint (Eq. 7) with Δ*µ* = 0. If the system is instead dissipative, with Δ*µ* ≠ 0, the shape of the flux ratio curve can change relative to the equilibrium system (SI Figure 1).

### Thermodynamic Basis for Substrate Selective Inhibition

To better understand the connections between our results and those of Mallory et al., ^9^ we can consider the coarse-graining, where we cannot explicitly detect enzyme states bound to the inhibitor. In this case, we will be unable to differentiate between states E and EI, ER and ERI, and other pairs of states for more complicated kinetic schemes. In the corresponding coarse-grained model, the pairs of states are lumped together to yield a simplified reaction scheme that can be easily analyzed.

For example, the competitive inhibition scheme (Figure 1A) can be coarse-grained to yield the effective reaction scheme

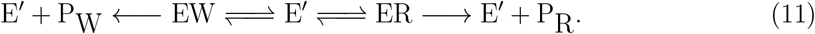

Here, we assume that the free enzyme (E) and the inhibited enzyme (EI) are indistinguishable, and are thus represented by the composite state (E′) so that

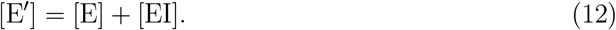

At steady-state, we have

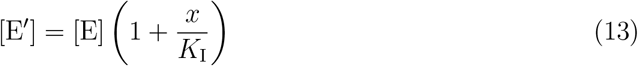

where *K*_I_ is the association constant for inhibitor binding,

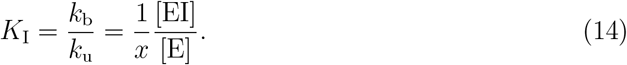

The effective substrate binding rates for the coarse-grained network are obtained by renormalizing the original rates. By re-normalizing the rates, the steady-state fluxes for each reaction in the coarse-grained network are identical to those in the original network. The effective substrate association rates for the coarse-grained network are given by

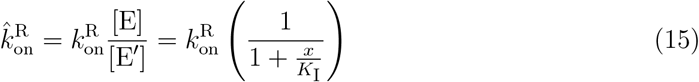

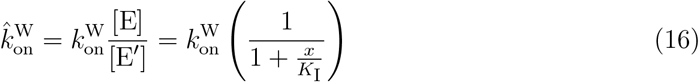

Here, increasing the inhibitor concentration effectively decreases the substrate association rate by limiting the concentration of free enzyme; this effect can be observed in the corresponding free-energy landscape, in which the presence of inhibitor lowers the effective energy of the coarse-grained free enzyme state (SI Figure 2A). The enzyme-substrate dissociation rates 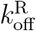 and 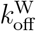 are unaffected by the presence of inhibitor; thus, competitive inhibitors effectively perturb the free-energy of the composite state [E′], and therefore the invariance of the flux ratio holds.

The uncompetitive inhibition scheme (Figure 1B) can be coarse-grained in a similar manner. Here, we make a similar assumption that the enzyme-substrate complexes (ER and EW) cannot be distinguished from the enzyme-substrate-inhibitor complexes (ERI and EWI), which leads to the effective reaction scheme

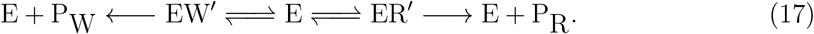

The effective substrate dissociation rates for the coarse-grained network are given by

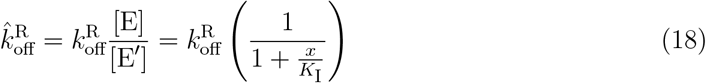

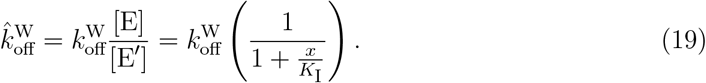

The effective catalytic rates for the coarse-grained network take a similar form and are given by

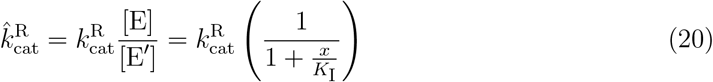

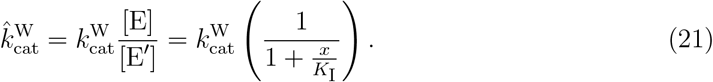

Changes in the inhibitor concentration effectively change the energies of the ER′ and EW′ states (SI Figure 2B), with the catalytic and dissociation rates being scaled by the same factor. The presence of the inhibitor does not affect the effective transition barriers for either substrate binding or catalysis; therefore, the flux ratio remains unaffected.

In the case of partial and mixed inhibition (Figure 1C,D), the inhibitor can bind to both the free enzyme (E) and the enzyme-substrate complex (ER and EW). Assuming again that the unbound and inhibitor-bound states cannot be distinguished, we can coarse-grain the network to yield the reduced scheme:

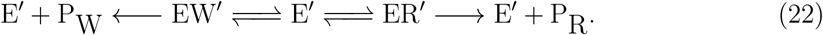

Below, we demonstrate that for this effective scheme, describing partial and mixed inhibition cases, changes in the inhibitor concentration alter the effective barrier heights in the coarsegrained networks.

For simplicity, we assume the catalytic rates 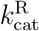 and 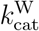 are much slower than the other transitions. In that case, we can make a quasi-equilibrium approximation to compute the effective substrate binding and unbinding rates. In that limit, for the coarse-grained network, the effective substrate binding rates are given by

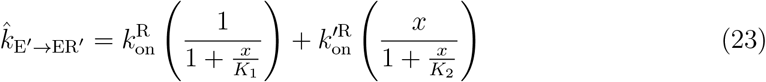

and

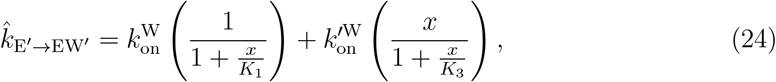

where *K*_1_, *K*_2_, and *K*_3_ are the association constants for inhibitor binding to the free enzyme and enzyme-substrate complexes (ER, EW), respectively.

Here, we see that the inhibitor affects the effective rate of both substrate binding and unbinding. By looking at the ratio of the effective binding rates for the R and W substrates

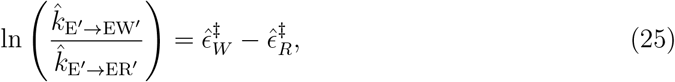

we can see how the inhibitor affects the relative free-energy barriers 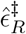 and 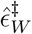 for the R and W substrates, and thus the underlying thermodynamic landscape. Using the coarse-grained network, the ratio is given by

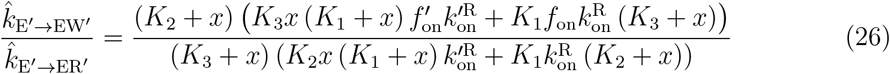

where each rate 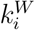 is redefined in terms of discrimination factors as 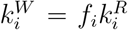. We see that the ratio is a function of the inhibitor concentration *x*, which implies, consistent with the results of Mallory et al., ^9^ that the inhibitor effectively acts as a perturbation affecting the relative free-energy barriers for the substrate binding reactions. Trivially, we can see that the inhibitor will be unable to affect the ratio of effective rates if both *K*_2_ = *K*_3_ and 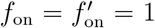; the ratio will deviate from unity for some inhibitor concentration *x* if either constraint is relaxed. Ultimately, we require either reaction cycles involving substrate binding or the existence of multiple pathways to the product formation reaction(s) for the inhibitor to affect the ratio of effective transition rates for reactions involving different substrates, and consequently, the difference in effective free-energy barriers for said reactions.

### Mixed and Partial Inhibition Can Yield Non-Monotonic Responses

Thus far, we have shown that mixed and partial inhibitors can affect enzymatic substrate selectivity. Beyond simply affecting substrate specificity, for some parameter regimes, it is possible to observe non-monotonic responses to changes in the inhibitor concentration. In these cases, the product formation flux ratio *η* can exceed the limits for the uninhibited enzyme (*η*_0_) and the enzyme fully saturated with inhibitor (*η*_∞_).

Two distinct classes of non-monotonic behaviors are possible depending on the particular model parameters; for simplicity, we constrain the parameters to ensure that the flux ratio at saturating concentration of inhibitor, *η*_∞_, is lower than that of the uninhibited enzyme, *η*_0_. This does not affect the generality of our conclusions because we can always redefine the right substrate as the one with decreasing flux. Non-monotonic behavior of *η* will imply non-monotonic behavior of 1/*η* and vice versa. First, for some parameterizations, an undershoot condition is observed, in which the substrate selection error can be improved beyond the saturating (η < η_∞_) limit for intermediate inhibitor concentrations (Figure 3A). Second, other parameterizations can display an overshoot, whereby the substrate selection error is higher than the uninhibited enzyme (*η* > *η*_0_) for some intermediate inhibitor concentrations (Figure 3B). Beyond undershoot and overshoot, in other cases, the substrate selection error is less sensitive to changes in inhibitor concentration at sub-saturating concentrations, yielding a plateau-like effect (SI Figure 3).

**Figure 3.**
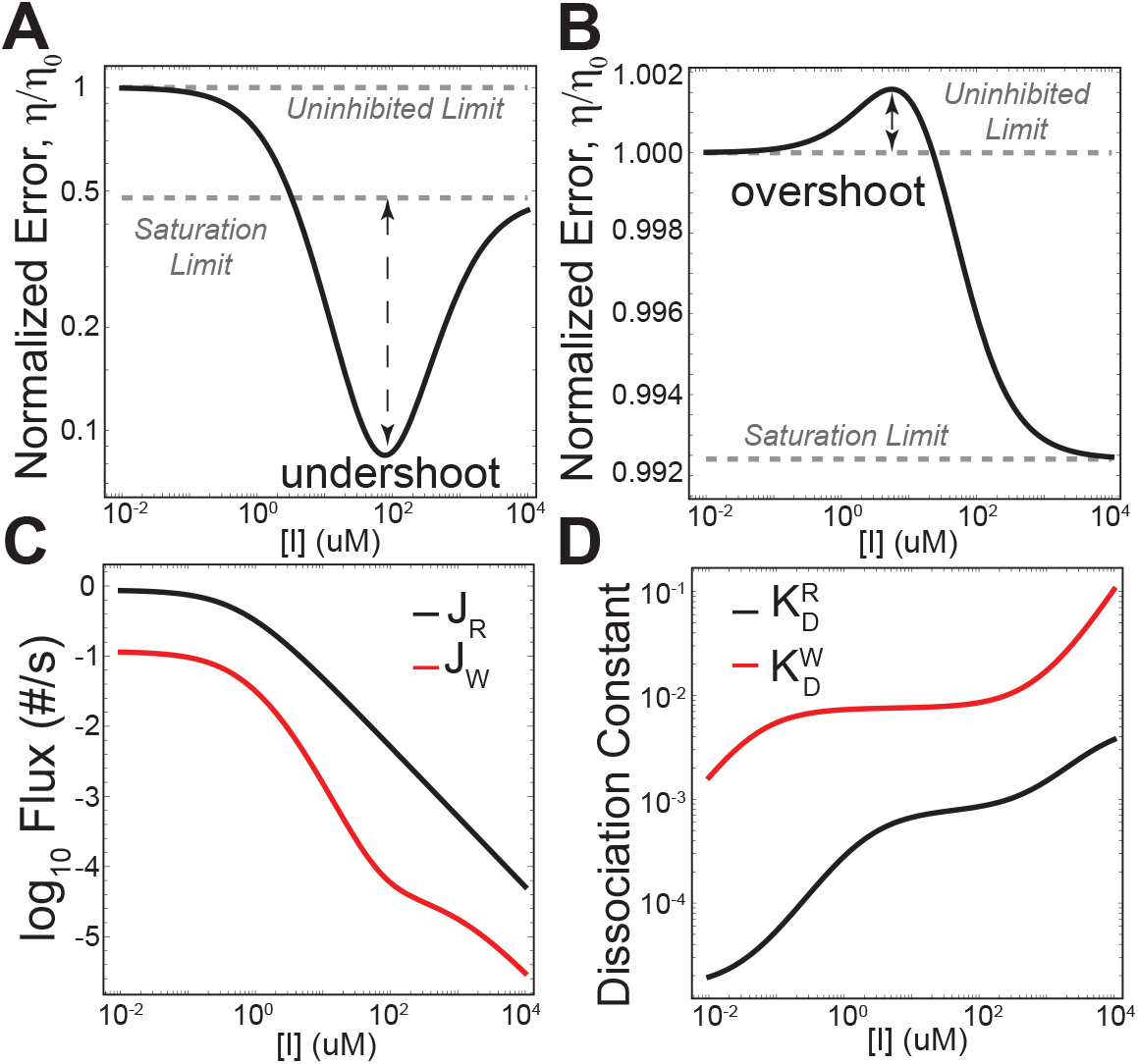
Substrate-selective enzyme inhibitors can, in principle, have non-monotonic effects on product flux ratios, including undershoot (A) and overshoot (B), where the flux ratio can exceed the limiting bounds. The non-monotonic response to inhibitor arises due to differential effects on the product formation fluxes (C) caused by changes in the effective substrate dissociation constants (D). The flux and effective dissociation constant curves shown correspond to the model in (A).

The uninhibited limit *η*_0_ for both mixed and partial inhibition is identical and corresponds to that of a basic Michaelis-Menten enzyme with two possible substrates, ^14^ which is

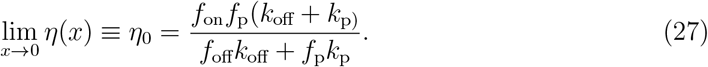

The expression for specificity is defined in terms of the rates for reactions involving the desired substrate, R, as well as discrimination factors *f*_*i*_ (Eq. 3).

In contrast, the saturation limit *η*_∞_ varies between mixed and partial inhibition. For mixed and non-competitive inhibition, the saturating limit is simply

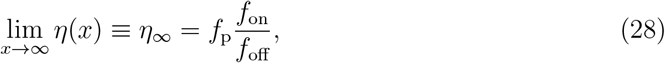

which is the discrimination in catalysis (*f*_p_) scaled by the ratio of discrimination in sub-strate binding and unbinding. The corresponding limit for partial inhibition is somewhat more complicated; because the enzyme retains residual catalytic activity when bound to the inhibitor, the saturating limit for partial inhibition takes the same structural form as for a typical Michaelis-Menten enzyme (Eq. 27), but with kinetic rates and discrimination factors that in principle may differ from the uninhibited enzyme:

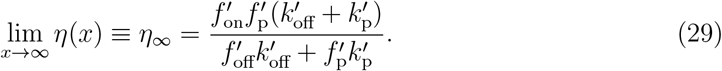

The underlying basis for the non-monotonic responses observed for mixed and partial inhibition can be probed by examining the changes in the product formation fluxes *J*_R_ and *J*_W_ and the effective substrate binding and unbinding rates as a function of inhibitor concentration. For one parameterization of the mixed inhibition model (Figure 1C) leading to undershoot (Figure 3A), the *J*_R_ and *J*_W_ fluxes become less sensitive to changes in inhibitor at different inhibitor concentration ranges, which causes the flux ratio to change non-monotonically as a function of inhibitor (Figure 3C). Ultimately, the effective substrate dissociation constant 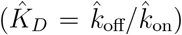 provides insights into how changes in inhibitor concentration can differentially affect the R and W product formation fluxes. As the inhibitor concentration increases, the effective dissociation constant for the R substrate saturates at a lower inhibitor concentration than the W substrate, which allows the product formation fluxes to change at different rates (Figure 3D). In terms of model parameters, one possible way to achieve this non-monotonicity is by varying the discrimination factors for substrate unbinding (*f*_off_) and catalysis (*f*_p_); with all other parameters held constant, changing these two discrimination factors can yield both undershoot and overshoot (SI Figure 4).

**Figure 4.**
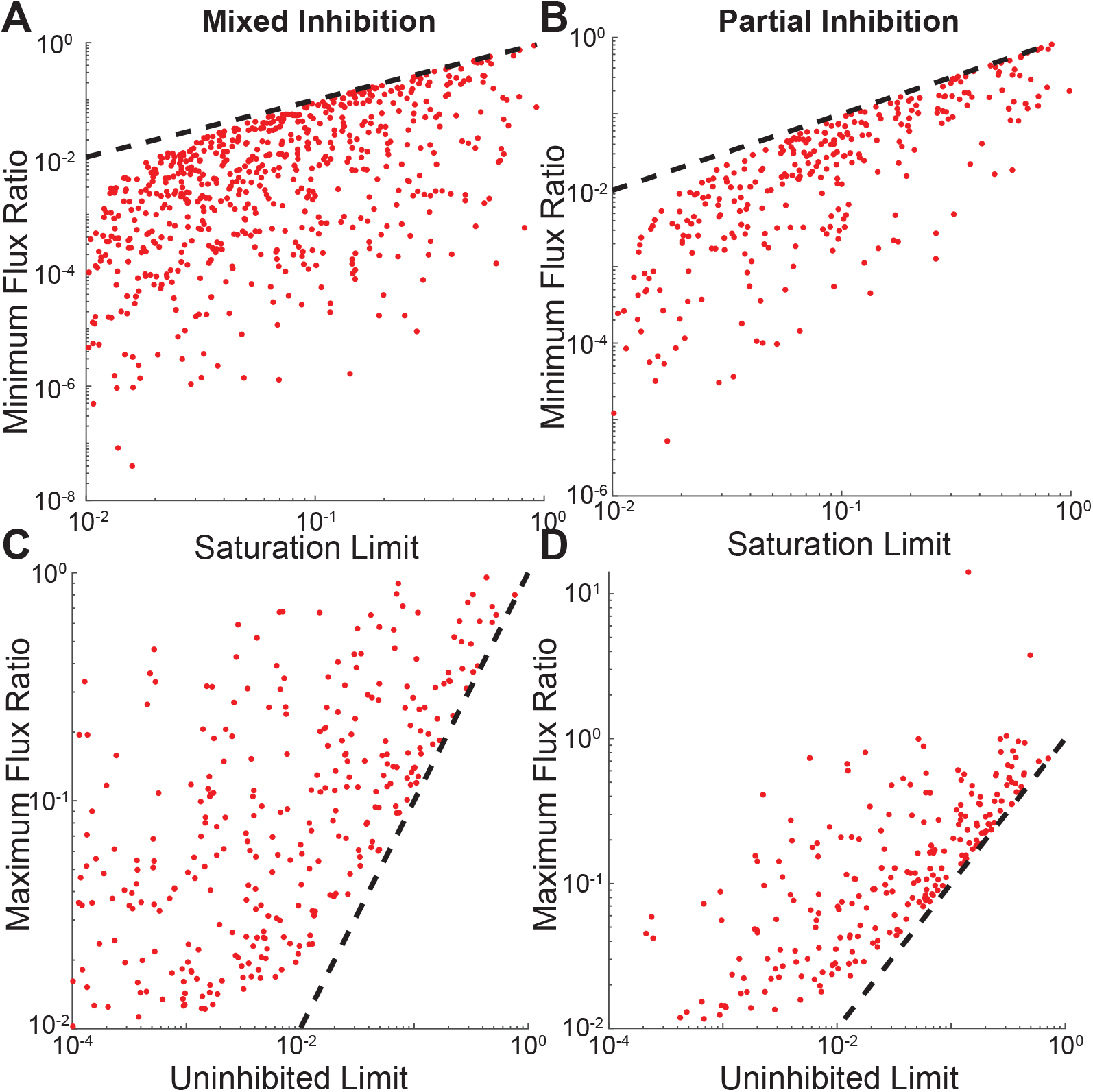
Mixed and partial inhibition can display both undershoot (A,B) and overshoot (C,D), where the flux ratio *η* can exceed the limiting bounds (*N* = 10000 parameter sets). The dashed lines correspond to points where either the minimum flux ratio is equal to the saturation limit or the maximum flux ratio is equal to the uninhibited limit; consequently, parameter sets that fall along these lines display neither overshoot nor undershoot. Parameter sets displaying only slight undershoot and overshoot, i.e. *q*_under_ (Eq. 8 and *q*_over_ (Eq. 9) less than *ϵ* = 0.01 are not plotted.

In order to verify that these non-monotonic behaviors are general, we randomly sampled 10000 models and checked for overshoot and undershoot numerically. We enforce the inequality

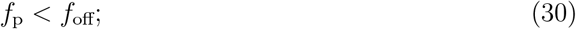

to ensure that the constraint *η*_∞_ < *η*_0_ is satisfied for the sampled parameters, Thus, as long as the inequality holds, the flux ratio at saturating inhibitor concentrations should be less than the uninhibited limit for mixed inhibitors. The situation for partial inhibition is complicated by the capacity for catalysis when bound to the inhibitor; therefore, when sampling parameters for both models, we reject parameter sets that lead to *η*_∞_ > *η*_0_. Under this sampling regime, we find that parameterizations yielding both undershoot (Figure 4A,B) and overshoot (Figure 4C,D) responses can be readily found for both mixed and partial inhibition modes. Interestingly, in practice, we find that, for randomly sampled parameter sets, the mixed and partial inhibition modes have different fractions of parameters that display undershoot and overshoot (SI Figure 5)

**Figure 5.**
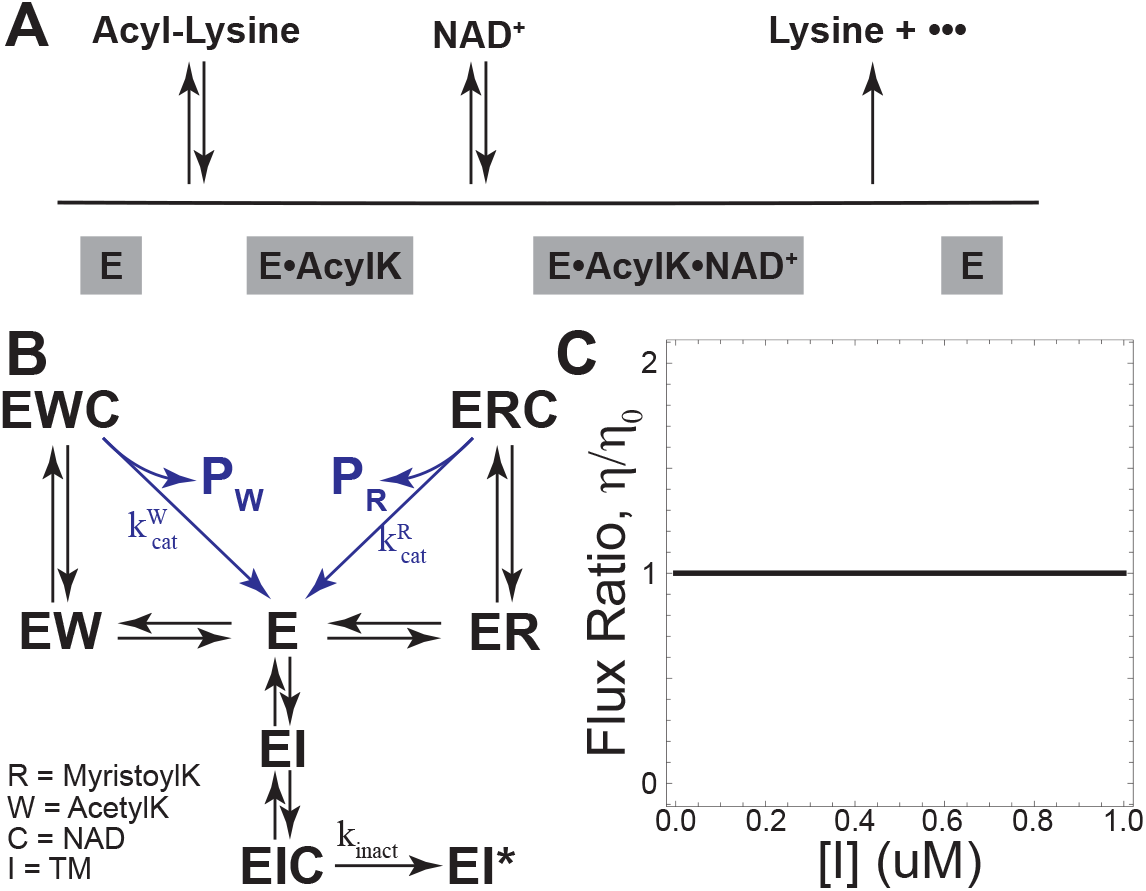
The NAD-dependent de-acylase SIRT2 is proposed to have an ordered binding mechanism, with acyl-lysine substrates binding prior to NAD (A, scheme adapted from^19^). If TM is competitive with the acyl-lysine substrate and uncompetitive with NAD, SIRT2 deacylation can be modeled with a simple kinetic scheme (B). Under these constraints, we predict that TM cannot affect SIRT2 substrate selectivity (C).

### Suicide Inhibition Cannot in Itself Affect Specificity

In addition to reversible inhibitors, mechanism-based inhibitors (also known as “suicide” inhibitors) are capable of permanently inactivating the enzyme. Such inhibitors compete with *bona fide* substrates for access to the active site in a manner analogous to competitive inhibitors, with a basic reaction scheme given by

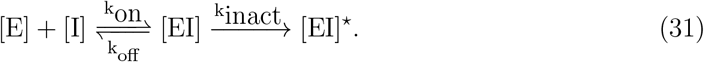

However, unlike real substrates, which can proceed through the full enzyme catalytic cycle, the suicide substrate becomes stuck at some covalent intermediate state, leading to enzyme inactivation ([EI]^⋆^, SI Figure 6A).

**Figure 6.**
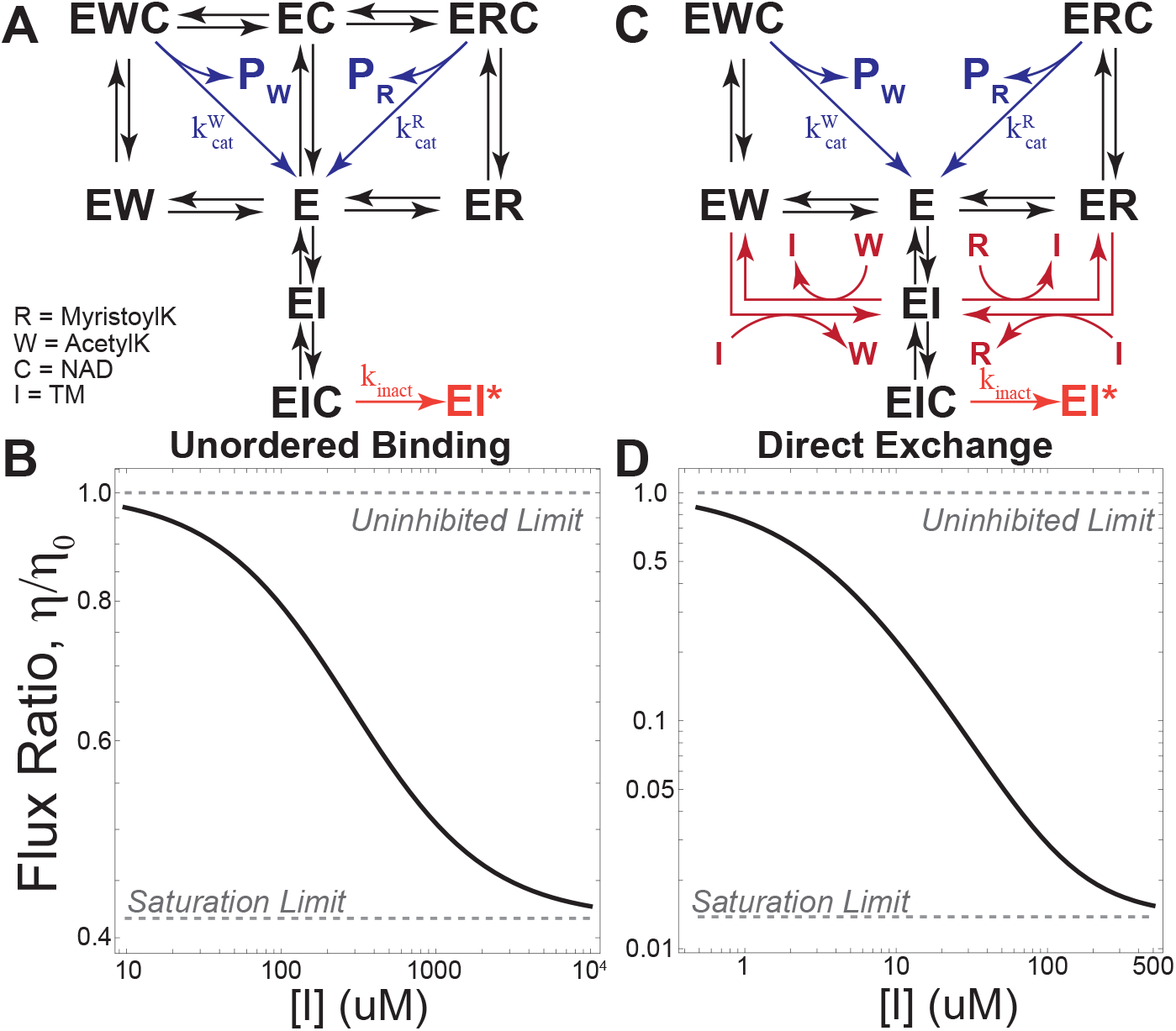
If the TM inhibitor is competitive with substrate and uncompetitive with NAD, either (A) unordered binding or (B) direct substrate exchange reactions are required for TM to affect substrate specificity (C,D).

At steady-state, for example, in cells, the quantity of active enzyme in the system remains constant over time; as enzyme is removed by inactivation, new enzyme is introduced into the system to maintain the steady state. In order to account for this, we modify the governing equation (Eq. 1) to include a production term *σ*(*x*) that can balance the loss of enzyme by inactivation, i.e.

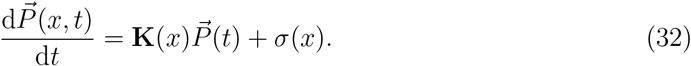

With this modification, we find that enzyme inactivation by suicide substrates cannot affect substrate specificity at steady state (SI Figure 6B). Additionally, time-course simulations of the suicide inhibition model rapidly approach the steady-state limit (SI Figure 6C). Further-more, even in the absence of enzyme replacement, the system remains largely unaffected by the addition of the inhibitor assuming fixed substrate concentrations (i.e. [R], [W] ¿¿ [E]); with inhibitor concentrations much higher than enzyme, the flux ratio exhibits a transient change upon inhibitor addition but quickly returns to the steady-state limit (SI Figure 7A,B).

Under conditions in which the inhibitor concentration is much lower than the concentration of substrate and enzyme, transient changes in substrate selectivity are minimal (SI Figure 7C).

### Ordered Binding Prevents TM from Affecting SIRT2 Specificity

The principles formulated for the basic inhibition networks can be applied to more complex schemes. Thus, we next applied our framework to a real system (Figure 5A). The Sirtuin family of NAD-dependent de-acylases is particularly interesting, as they are involved in many biological processes, including apoptosis and inflammation;^15^ in addition, SIRT2 has been implicated in some cancers.^16^ Recently, a thiomyristoyl lysine compound (TM) was developed as a mechanism-based inhibitor of the Sirtuin family member SIRT2;^17^ like other suicide inhibitors, TM proceeds down the SIRT2 catalytic pathway but stalls at an intermediate step and remains covalently linked to the enzyme. Notably, TM has been shown to inhibit SIRT2 histone 3 lysine 9 (H3K9) deacetylation while leaving SIRT2 activity towards other substrates, such as myristoylated peptides, less affected. ^7,17^

The precise mechanism by which TM affects SIRT2 substrate selectivity is unclear. It has been proposed that the effect of TM on SIRT2 substrate specificity can be explained as a function of relative substrate binding affinities; ^7^ if the inhibitor binds SIRT2 with a binding affinity intermediate between some desired “correct” substrate and an undesired “incorrect” substrate, it may be capable of preferentially inhibiting SIRT2 action against the incorrect substrate.

Kinetic analyses by Jing et al. provided insights into the TM inhibition mechanism. We employ these results to develop a model of SIRT2 inhibition. Competition assays under either NAD saturating conditions (1000 nM NAD) or (2) substrate saturating conditions (100 nM acetylated H3K9) suggest that TM is competitive with substrate and uncompetitive with NAD.^17^ Additionally, SIRT2 has been reported to follow an ordered binding mechanism; ^18,19^ with ordered binding, the acyl lysine substrate must bind to SIRT2 to enable NAD binding.

With these constraints, we arrive at the model in Figure 5B. Lineweaver-Burke (LB) plots generated with the ordered binding model still predict substrate-competitive inhibition by TM (SI Figure 8A,B), consistent with the results of Jing et al.

Notably, the ordered binding model lacks reaction cycles involving the possible substrates. Thus, we expect SIRT2 substrate selectivity to be insensitive to TM concentration. Indeed, we find 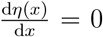 for all possible parameters, and that TM cannot affect SIRT2 specificity while still satisfying the constraints imposed by experimental evidence (Figure 5C).

### Substrate-Selective Inhibition of SIRT2 Requires Direct Substrate Exchanges or Unordered Binding

In order for TM to affect SIRT2 substrate selectivity, we must introduce additional pathways leading to product formation into the network in Figure 5A. There are at least two ways this can be achieved without changing our assumptions on the mode of TM inhibition. First, we can abandon the unordered binding constraint, and allow SIRT2 to bind NAD and acyl lysine substrates in a random order (Figure 6A); as expected, relaxing the ordered binding constraint allows TM to affect SIRT2 substrate specificity (Figure 6B). Additionally, LB plots generated from the unordered binding model still largely agree with a substrate-competitive inhibition mechanism (SI Figure 8C,D).

Alternatively, we can introduce direct substrate exchange reactions. In such reactions, either substrate binding or inhibitor binding can displace the currently bound substrate, which leads to the network in Figure 6C. As with relaxing the ordered binding constraint, introducing direct exchange reactions has the effect of creating cycles of reactions within the network; furthermore, with direct exchange reactions there is no need to relax the ordered binding constraint. Again, we see that allowing for direct exchanges enables TM to affect SIRT2 substrate specificity (Figure 6D).

Similar to the unordered binding model, the direct exchange model again agrees with substrate-competitive inhibition by TM (SI Figure 8E,F). However, at low values of 1/[R], the model predicts a slight curving in the LB plot; this curving was also observed in some cases for the unordered binding model. The curving may be an artifact of the precise parameters chosen for the model, which are largely unknown for SIRT2. However, if the effect is real, it could possibly be used to differentiate between the unordered binding and direct exchange models as long as the curving is significant; if the curving is too subtle, it could be missed as experimental noise.

## Discussion

Here we have presented an analysis of elementary enzyme inhibition schemes for their capacity to influence enzyme substrate selectivity. Through mathematical modeling, we have shown that competitive and uncompetitive inhibition schemes are fundamentally incapable of affecting substrate selectivity (Figure 2). Previous experimental studies appear to corroborate this result for competitive inhibition, where enzyme active site inhibitors have generally failed to affect substrate selectivity. ^6^

The ability of non-competitive and mixed inhibitors, as well as partial inhibitors, to affect substrate selectivity results from the presence of reaction cycles within the respective governing kinetic schemes. In these systems, cycles of reactions (e.g. loops) allow the inhibitor to perturb the effective transition free-energy barrier for transitions between states involving the substrate (such as substrate binding), and consequently, affect enzyme substrate selectivity in a manner consistent with the results of Mallory et al. ^9^ However, reaction cycles must involve both substrate and inhibitor binding to affect substrate selectivity; in a modified version of the SIRT2 ordered binding model where SIRT2 binds to NAD prior to the substrate (a mechanism similar to the related SIRT6^20^), the reaction cycle involves only NAD. Under this alternative reaction mechanism, TM cannot affect specificity (SI Figure 9).

Notably, while reaction cycles that involve both inhibitor and substrate binding are sufficient for substrate selectivity, they are not a strict requirement. In some systems, such as the inhibited IDE enzyme studied by Maianti et al., ^6^ multiple catalytically competent states exist for one or more possible substrates even when bound to inhibitor. In the case of IDE, the proposed kinetic scheme is essentially a limiting case of the partial inhibition scheme (SI Figure 10A, scheme adapted from^6^), whereby inhibitor is incapable of binding to the substrate-bound enzyme; under this scheme, no reaction cycles are present. However, the existence of multiple catalytically-competent enzyme states (in this case the ER and ERI states for the R substrate) allows substrate-selective inhibition to be maintained even in the absence of reaction cycles (SI Figure 10B). With multiple catalytic states for one or both substrates, the overall product formation flux ratio is a ratio of sums of fluxes and can therefore change as a function of inhibitor concentration as inhibitor-bound catalytic states become more prevalent.

Finally, in our models, we have considered only the average behavior of inhibited enzymes. However, Robin et al. have demonstrated that inhibitor binding can paradoxically increase enzyme turnover rates (catalytic events per unit time) at intermediate concentrations, as measured at the single-enzyme level.^21^ In their framework, they abandon the typical assumption that system state transitions are Markovian and are exponentially distributed and instead allow the distribution of transition waiting times to follow an arbitrary distribution. While our models predict that competitive and uncompetitive inhibition are incapable of affecting specificity at the bulk enzyme level (Figure 2A,B), it is possible that these inhibition modes might show some effect on specificity at the single-enzyme level. Future work could extend our models in an analogous manner to test whether these effects can affect substrate selectivity.

## Conclusions

Overall, we have shown that mixed and partial inhibition are the only reversible inhibition mechanisms capable of affecting substrate selectivity. The physical basis of this selectivity can be explained by analogy to the work of Mallory et al., ^9^ which demonstrated that perturbations in the free-energy barriers between states can affect biochemical pathway flux ratios. In mixed and partial inhibition schemes, as well as in systems possessing reaction cycles, the presence of an inhibitor (or other biochemical cofactors) acts as a *de facto* perturbation, affecting the transition barriers. Changes in substrate selectivity, which are nothing more than changes in ratios of biochemical pathway fluxes, are a direct consequence of these perturbations.

We have also demonstrated that the inhibitor TM, targeting the enzyme SIRT2, cannot affect substrate specificity while satisfying known biochemical constraints. The SIRT2 or-dered binding mechanism in which substrate binds prior to the required NAD cofactor,^19^ coupled with competitive inhibition of substrate binding, has the effect of eliminating reaction cycles that involve the acylated substrates; as a consequence, the inhibitor is not capable of affecting the transition barriers, and thus either a relaxation of constraints or the introduction of direct substrate exchange reactions is required to enable substrate-selective inhibition.

## Supporting information

Supporting Information Combined

## Acknowledgement

The authors would like to thank Anatoly B. Kolomeisky, Seokjoo Chae, and Zheng Diao for insightful comments on the manuscript.

O.A.I. acknowledges funding support from the Welch Foundation (Grant C-1995).

## Supporting Information Available

Detailed description of the reversible and suicide inhibition models; Effects of dissipation on flux ratios; coarse-grained free-energy landscape for competitive and uncompetitive inhibition; example of additional non-monotonic behavior; example of parameter regime yielding non-monotonic behavior; fraction of sampled models displaying non-monotonic behaviors; simulations of suicide inhibition model; Lineweaver-Burke plots for SIRT2 model variants; simulation of alternative SIRT2 ordered binding mechanism; simulation of Maianti IDE model; table of sampling bounds; table of suicide inhibition model parameters (PDF)

