## Supporting Information Combined for "Effects of chemical modulators on enzyme specificity"

### Supplemental Methods

#### Model Implementation

The inhibition models are implemented in terms of the forward chemical master equation, which is defined by a system of ordinary differential equations

$$\frac{\mathrm{d}\vec{P}(t)}{\mathrm{d}t} = \mathbf{K}\vec{P}(t) + \vec{\sigma}(t) \quad (1)$$

with the constraint

$$\sum_i P_i(t) = 1 \quad (2)$$

to ensure normalized probabilities. The matrix $\mathbf{K}$ encodes the state transitions for each system, with $\mathbf{K}_{ji} = k_{ij}$ for each transition $i \to j$. The vector $\vec{\sigma}(t)$ ensures that the system is balanced such that the total amount of enzyme does not change in time, with

$$\sigma_i = \begin{cases} \sum_j J_{ji} & i = 1 \\ 0 & i \neq 1 \end{cases}$$

where each $J_{ji}$ is a flux from state $j$ to state $i$.

With the additional constraint

$$\frac{\mathrm{d}\vec{P}(t)}{\mathrm{d}t} = [0, 0, \cdots, 0]^{\mathrm{T}} \quad (3)$$

we can compute the steady-state probabilities for each system. The ratio of product formation fluxes is then readily computed; for a system with two possible substrates R and W, we can define the flux ratio as a function of inhibitor concentration as

$$\eta(x) = J_{\mathrm{W}}(x)/J_{\mathrm{R}}(x). \quad (4)$$

### Competitive Inhibition Model

The matrix defining the competitive inhibition model is given by:

$$\mathbf{K} = \begin{pmatrix} -xk_b - f_{\text{on}}k_{\text{on}} - k_{\text{on}} & k_{\text{off}} & f_{\text{off}}k_{\text{off}} & k_u \\ k_{\text{on}} & -k_{\text{off}} - k_p & 0 & 0 \\ f_{\text{on}}k_{\text{on}} & 0 & -f_{\text{off}}k_{\text{off}} - f_p k_p & 0 \\ xk_b & 0 & 0 & -k_u \end{pmatrix}$$

### Uncompetitive Inhibition Model

The matrix defining the uncompetitive inhibition model is given by

$$\mathbf{K} = \begin{pmatrix} -f_{\text{on}}k_{\text{on}} - k_{\text{on}} & k_{\text{off}} & f_{\text{off}}k_{\text{off}} & 0 & 0 \\ k_{\text{on}} & -xk_b - k_{\text{off}} - k_p & 0 & k_u & 0 \\ f_{\text{on}}k_{\text{on}} & 0 & -xf_bk_b - f_{\text{off}}k_{\text{off}} - f_pk_p & 0 & f_uk_u \\ 0 & xk_b & 0 & -k_u & 0 \\ 0 & 0 & xf_bk_b & 0 & -f_uk_u \end{pmatrix}$$

### Mixed Inhibition Model

The matrix defining the mixed inhibition model is given by:

$$\mathbf{K} = \begin{pmatrix} -xk_b - f_{\text{on}}k_{\text{on}} - k_{\text{on}} & k_{\text{off}} & k_u & 0 & f_{\text{off}}k_{\text{off}} & 0 \\ k_{\text{on}} & -xk'_b - k_{\text{off}} - k_p & 0 & k'_u & 0 & 0 \\ xk_b & 0 & -f'_{\text{on}}k'_{\text{on}} - k'_{\text{on}} - k_u & k'_{\text{off}} & 0 & f'_{\text{off}}k'_{\text{off}} \\ 0 & xk'_b & k'_{\text{on}} & -k'_{\text{off}} - k'_u & 0 & 0 \\ f_{\text{on}}k_{\text{on}} & 0 & 0 & 0 & -xf'_bk'_b - f_{\text{off}}k_{\text{off}} - f_pk_p & f'_uk'_u \\ 0 & 0 & f'_{\text{on}}k'_{\text{on}} & 0 & xf'_bk'_b & -f'_{\text{off}}k'_{\text{off}} - f'_uk'_u \end{pmatrix}$$

### Partial Inhibition Model

The matrix defining the partial inhibition model is given by:

$$\mathbf{K} = \begin{pmatrix} -xk_b - f_{\text{on}}k_{\text{on}} - k_{\text{on}} & k_{\text{off}} & k_u & 0 & f_{\text{off}}k_{\text{off}} & 0 \\ k_{\text{on}} & -xk'_b - k_{\text{off}} - k_p & 0 & k'_u & 0 & 0 \\ xk_b & 0 & -f'_{\text{on}}k'_{\text{on}} - k'_{\text{on}} - k_u & k'_{\text{off}} & 0 & f'_{\text{off}}k'_{\text{off}} \\ 0 & xk'_b & k'_{\text{on}} & -k'_{\text{off}} - k'_p - k'_u & 0 & 0 \\ f_{\text{on}}k_{\text{on}} & 0 & 0 & 0 & -xf'_b k'_b - f_{\text{off}}k_{\text{off}} - f_p k_p & f'_u k'_u \\ 0 & 0 & f'_{\text{on}}k'_{\text{on}} & 0 & xf'_b k'_b & -f'_{\text{off}}k'_{\text{off}} - f'_p k'_p - f'_u k'_u \end{pmatrix}$$

### Transient Suicide Inhibition Model

The suicide inhibition model is implemented as a system of ordinary differential equations (ODEs):

$$\frac{\mathrm{d}\,[\mathrm{E}]}{\mathrm{d}t} = \left(f_{\text{off}}k_{\text{off}}\,[\mathrm{EW}] + k_{\text{off}}\,[\mathrm{ER}] + k_{\text{u}}\,[\mathrm{EI}]\right) + \left(f_{\text{cat}}k_{\text{cat}}\,[\mathrm{EW}] + k_{\text{cat}}\,[\mathrm{ER}]\right) - \left(f_{\text{on}}k_{\text{on}}\,[\mathrm{W}] + k_{\text{on}}\,[\mathrm{R}] + k_{\text{b}}\,[\mathrm{I}]\right)[\mathrm{E}] \tag{5}$$

$$\frac{\mathrm{d}\,[\mathrm{I}]}{\mathrm{d}t} = k_{\text{u}}\,[\mathrm{EI}] - k_{\text{b}}\,[\mathrm{E}]\,[\mathrm{I}] \tag{6}$$

$$\frac{\mathrm{d}\,[\mathrm{ER}]}{\mathrm{d}t} = k_{\text{on}}\,[\mathrm{E}]\,[\mathrm{R}] - \left(k_{\text{off}} + k_{\text{cat}}\right)[\mathrm{ER}] \tag{7}$$

$$\frac{\mathrm{d}\,[\mathrm{EW}]}{\mathrm{d}t} = k_{\text{on}}\,[\mathrm{E}]\,[\mathrm{W}] - \left(f_{\text{off}}k_{\text{off}} + f_{\text{cat}}k_{\text{cat}}\right)[\mathrm{ER}] \tag{8}$$

$$\frac{\mathrm{d}\,[\mathrm{EI}]}{\mathrm{d}t} = k_{\text{b}}\,[\mathrm{E}]\,[\mathrm{I}] - \left(k_{\text{u}} + k_{\text{inact}}\right)[\mathrm{ER}] \tag{9}$$

The simulation was conducted with initial concentrations of enzyme and substrate equal to [E] = 0.2 uM, [R] = 200 uM, and [W] = 200 uM; the concentrations [R] and [W] of the two substrates are sufficiently large compared to the free enzyme [E] that the dynamics of the free substrate can be ignored. The model is first pre-equilibrated by integrating the system of ODEs in the absence of inhibitor ([I] = 0) with $k_{\text{cat}} = 0.275$ for 120 seconds. After pre-equilibration, the inhibitor concentration is increased to either [I] = 0.02 or [I] = 20 uM; the enzyme inactivation rate is set to $k_{\text{inact}} = 0.0275$. The pre-equilibrated model is then integrated again for 120 seconds to simulate the dynamics upon inhibitor addition. Other model rate constants are set according to the values in SI Table 2.

The product formation fluxes $J_{\text{R}}$ and $J_{\text{W}}$ are computed at each time-point by calculating

$J_{\mathrm{R}} = k_{\mathrm{cat}}\,[\mathrm{ER}]$ and $J_{\mathrm{W}} = f_{\mathrm{cat}} k_{\mathrm{cat}}\,[\mathrm{EW}]$. With the product formation fluxes, the flux ratio $\eta$ can be computed as

$$\eta = \frac{J_{\mathrm{W}}}{J_{\mathrm{R}}}. \tag{10}$$

In all cases, the system of ODEs was integrated using the built-in `ode89` solver in MATLAB® 2025b. The absolute and relative tolerances for the `ode89` solver were set to 1e-15 and 1e-12, respectively.

### Supplemental Figures

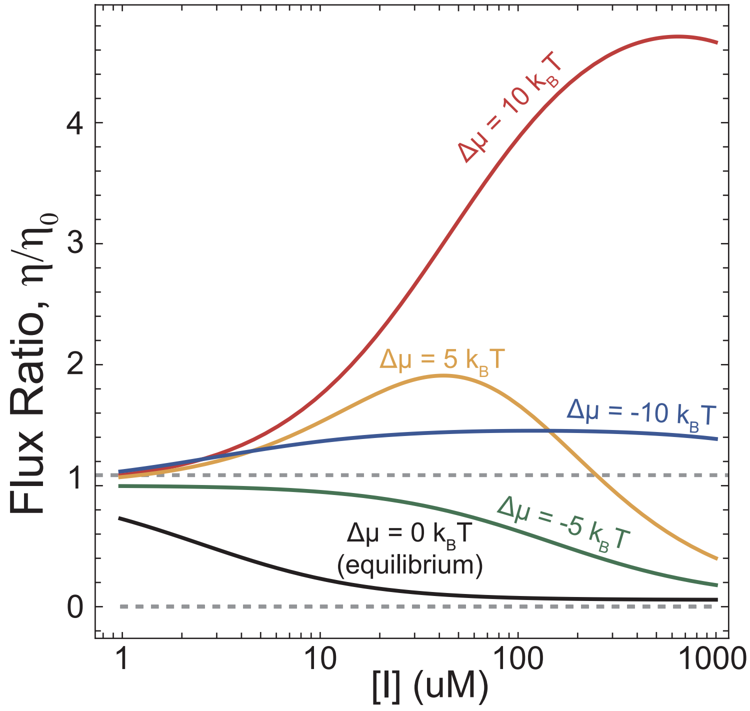


Figure S1: Dissipation can affect the response to increasing inhibitor concentrations. For the non-competitive model, dissipation can cause the flux ratio to exceed the equilibrium limits (gray dashed lines).

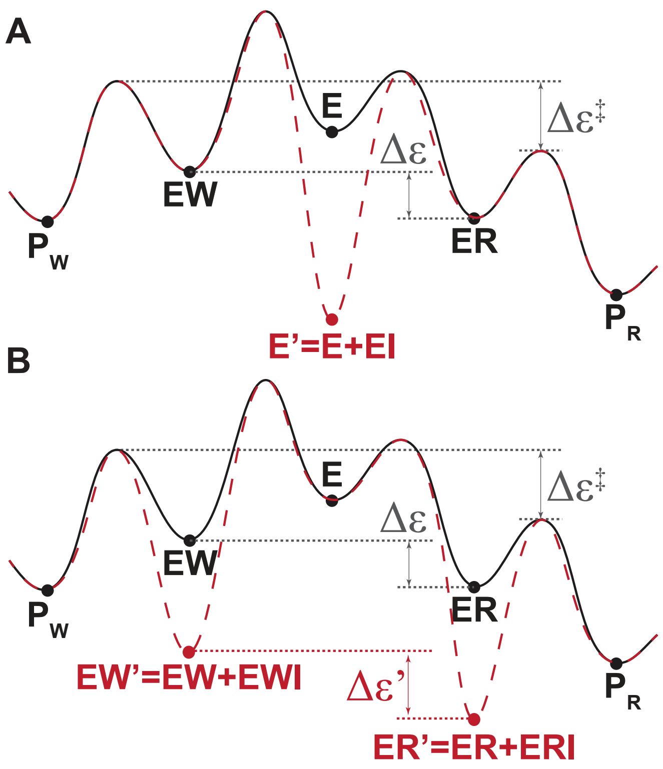


Figure S2: The coarse-grained energy landscape for the (A) competitive and (B) uncompetitive inhibitor suggests that the presence of inhibitor does not alter the effective difference in transition barrier heights ($\Delta\epsilon^\ddagger$) between the R and W substrates.

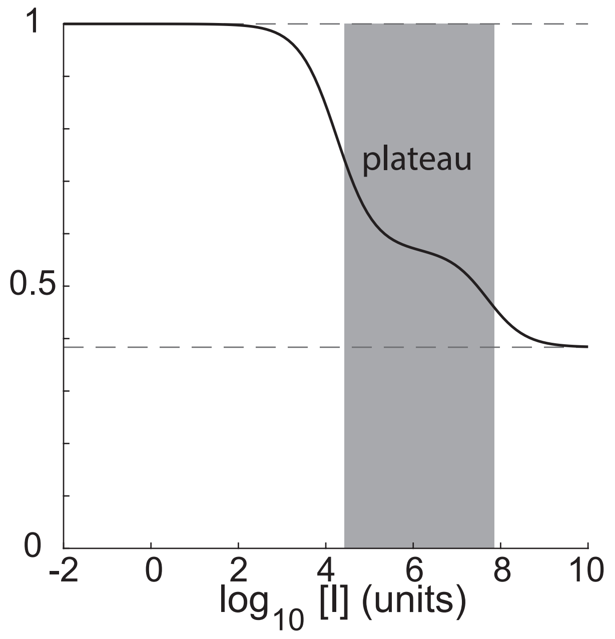


Figure S3: For some parameterizations, mixed and partial inhibition can display decreased sensitivity to changes in inhibitor concentration for intermediate concentration ranges.

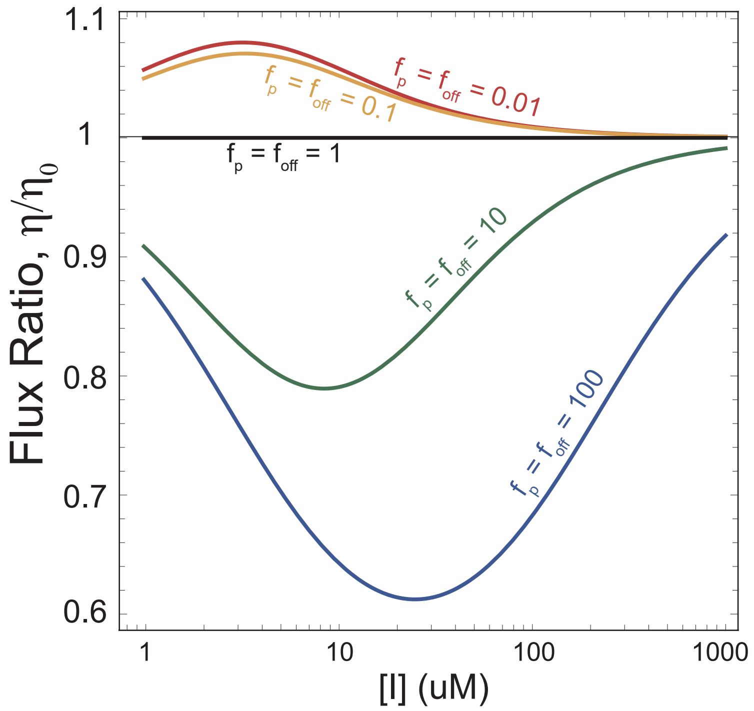


Figure S4: Varying the value of the parameters $f_\mathrm{p}$ and $f_\mathrm{off}$ appears to control the type of non-monotonic behavior observed for the mixed inhibition model. Here, $\mu = 0$ and all other parameters are set to unity.

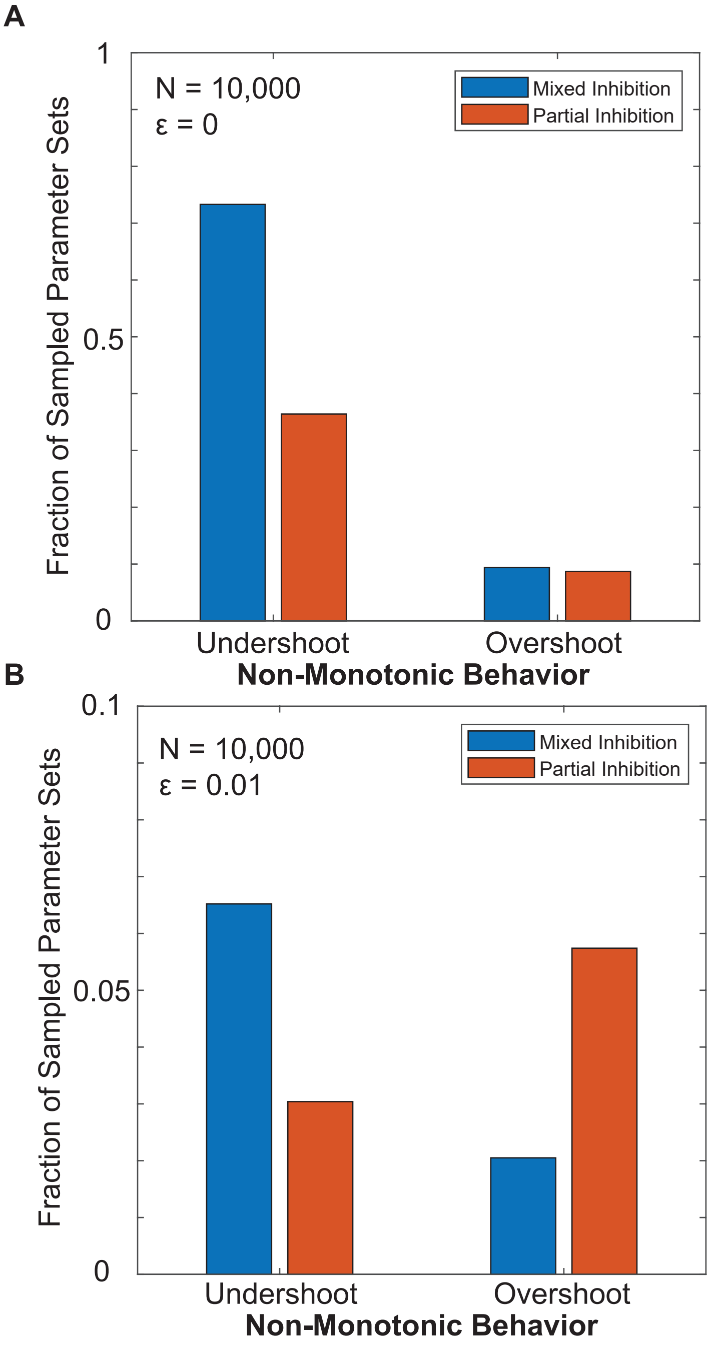


Figure S5: Mixed inhibition is generally more likely to display undershoot when the threshold $\epsilon$ is set to zero (A). In contrast, when the threshold $\epsilon = 0.01$, mixed and partial inhibition are more likely to display undershoot and overshoot, respectively (B).

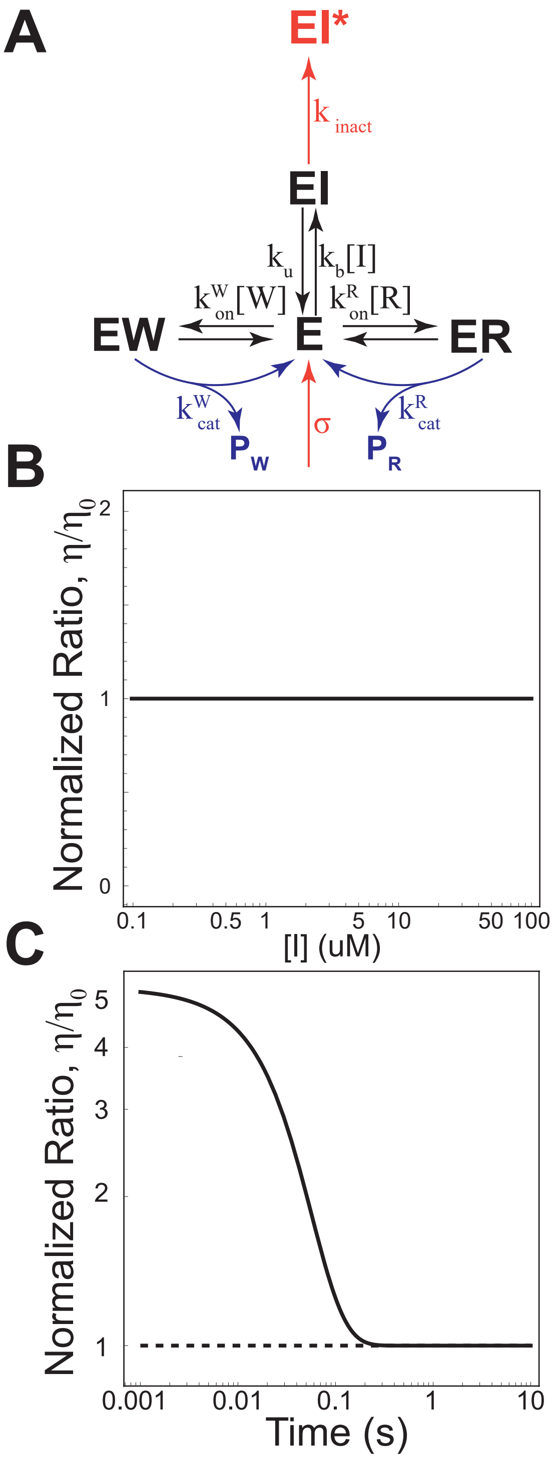


Figure S6: Mechanism-based inhibitors cannot undergo the full enzymatic cycle and cause the enzyme to become permanently inactivated (A); product formation reactions are shown in blue and enzyme inhibition due to inhibitor is shown in orange. At steady-state and replacement of inactivated enzyme (with rate $\sigma$), suicide inhibition cannot affect substrate specificity (B). Further, time-course simulations reveal that the system rapidly converges to the steady-state, uninhibited limit, $\eta/\eta_0 = 1$ (C).

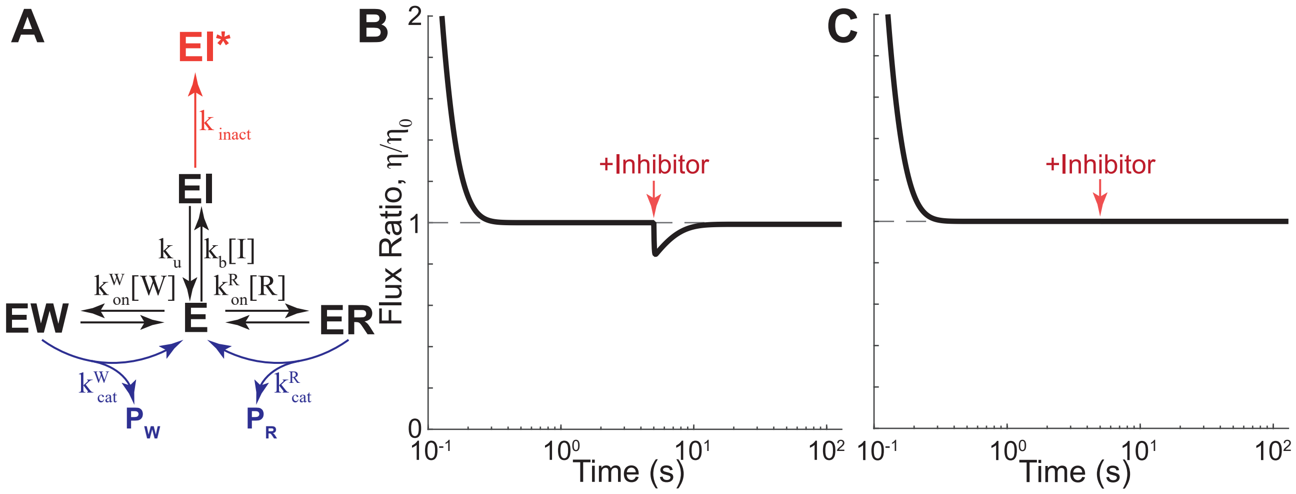


Figure S7: Suicide inhibition scheme without enzyme replacement (A). With constant substrate concentrations and inhibitor in great excess relative to the enzyme ([I] = 20 uM), the addition of a suicide inhibitor causes a transient change in substrate specificity (B). When the inhibitor concentration is much lower than that of the enzyme ([I] = 0.02 uM), the transient change is negligible (C). In both cases, the simulation is conducted with [I] = 0 uM at time t = 0 seconds, [R] = [W] = 100 uM, and [E] = 0.2 uM. Inhibitor is added at time t = 5 seconds and the simulation is then continued for an additional 120 seconds.

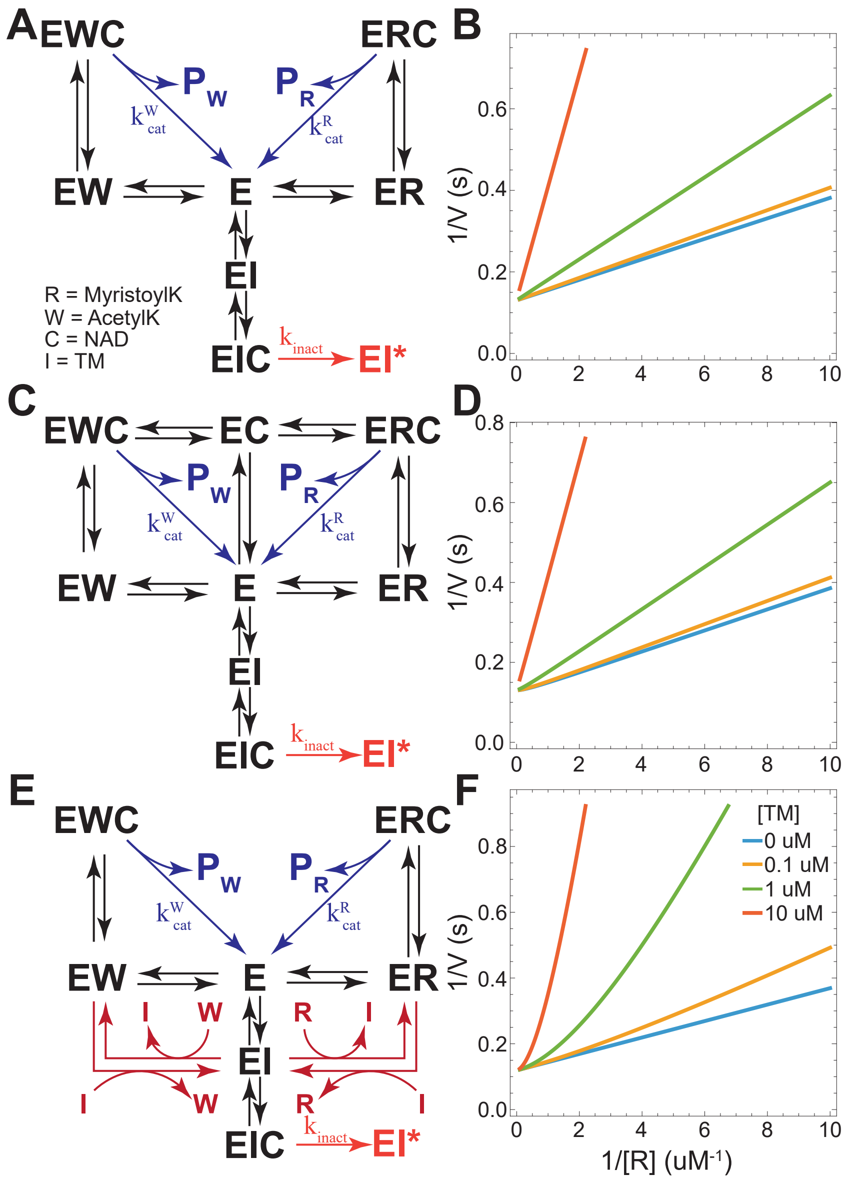


Figure S8: Lineweaver-Burke (LB) plots using the ordered binding SIRT2 model (A) are consistent with competitive inhibition by TM (B), and agree with the results of Jing et al. 2016.[1] Unordered substrate and NAD binding (C) additionally produces LB plots qualitatively consistent with experiment (D). With direct exchange reactions between substrates and inhibitor (E), the model is again largely consistent with experiment, but predicts slight curving in the LB plots at low values of 1/[R] (F).

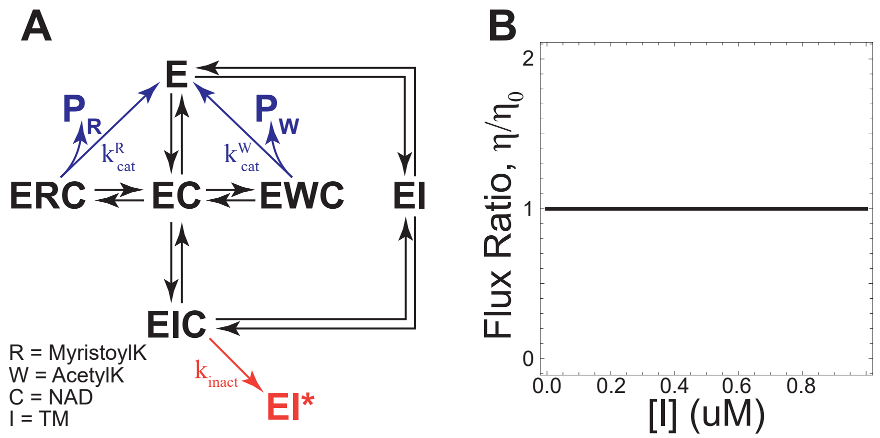


Figure S9: If SIRT2 followed an ordered binding mechanism in which NAD binds to the free enzyme prior to substrate, the only remaining thermodynamic cycle would involve NAD and TM (A). Under this mechanism, SIRT2 substrate selectivity cannot be affected by TM concentration (B). Product formation reactions are shown in blue and enzyme inhibition due to TM is shown in orange.

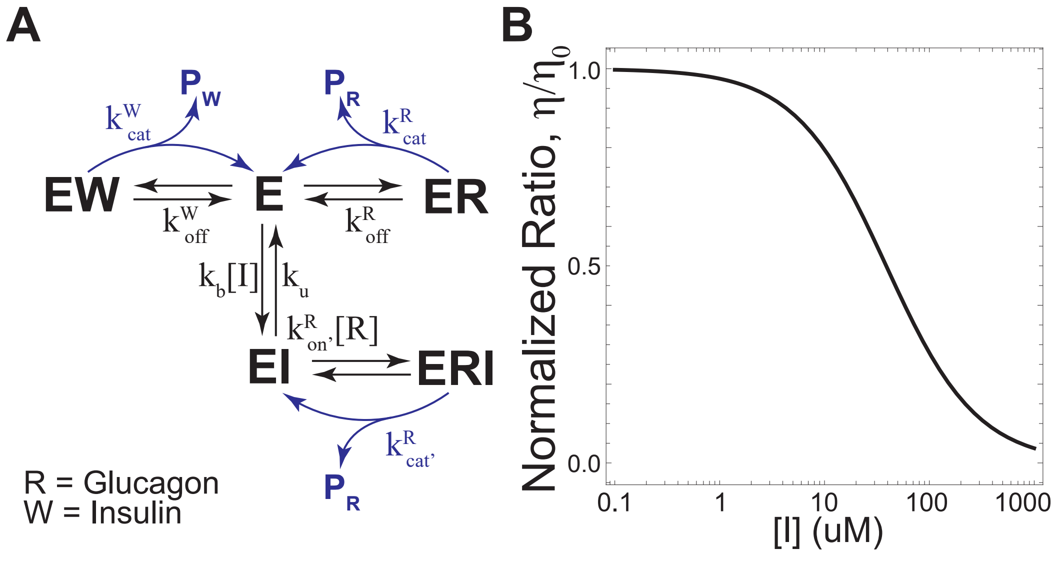


Figure S10: Maianti et al. propose that IDE can still bind and degrade glucagon when bound to a substrate-selective inhibitor, while blocking insulin degradation (A, scheme adapted from[2]). Under this scheme, there are no kinetic cycles, but the presence of two catalytically competent states for glucagon degradation (ER and ERI) compared to insulin (EW) enables substrate selective inhibition (B).

### Supplemental Tables

**Table S1: Discrimination Factor Sampling Bounds**

| Parameter | Lower Bound | Upper Bound |
|---|---|---|
| $f_{\text{on}}$ | $10^{-4}$ | $10^{0}$ |
| $f_{\text{off}}$ | $10^{0}$ | $10^{4}$ |
| $f'_{\text{b}}$ | $10^{-4}$ | $10^{0}$ |
| $f'_{\text{u}}$ | $10^{0}$ | $10^{4}$ |
| $f'_{\text{on}}$ | $10^{-4}$ | $10^{0}$ |
| $f'_{\text{off}}$ | $10^{0}$ | $10^{4}$ |

**Table S2: Suicide Inhibition Model Parameters**

| Parameter | Value | Units |
|---|---|---|
| $k_{\text{on}}$ | 40 | $\text{s}^{-1}\mu\text{M}^{-2}$ |
| $k_{\text{off}}$ | 0.5 | $\text{s}^{-1}$ |
| $k_{\text{b}}$ | 40 | $\text{s}^{-1}\mu\text{M}^{-2}$ |
| $k_{\text{u}}$ | 0.5 | $\text{s}^{-1}$ |
| $k_{\text{cat}}$ | 0.275 | $\text{s}^{-1}$ |
| $k_{\text{inact}}$ | 0.0275 | $\text{s}^{-1}$ |
| $f_{\text{on}}$ | 0.675 | unitless |
| $f_{\text{off}}$ | 94 | unitless |
| $f_{\text{cat}}$ | 0.0655 | unitless |
